# Sex Discordance and Risk of Breast Cancer, A Twin Study

**DOI:** 10.1101/621417

**Authors:** Livingstone Aduse-Poku, Shayesteh Jahanfar

## Abstract

**Objective:** The purpose of the study is to perform an analysis of the relationship between sex discordance and risk of breast cancer in female twins in the United States.

**Methods:** A cross-sectional study of 14,462 female twins was conducted using data from Washington State Twin Registry (WSTR) in the USA. The variables collected included, BMI, age, race and zygosity. This study used Generalized Estimating Equation (GEE) modeling to determine the relationships between twin pairs and variables of interest such as breast cancer and sex concordance. Zygosity, BMI, age and race were used for adjustment. Proband wise concordance was done to ascertain the heritability of breast cancer in twins.

**Results:** Being a female-female twin pair increased the odds of breast cancer by 34% (95%CI: 1.18-1.53). After adjusting for zygosity, age, BMI, race, and childbirth, the odds of breast decreased by 31% in female-female twin pairs [AOR (95%CI):0.69 (0.53-0.90)]. The proband wise concordance was higher in monozygotic twins as compared to dizygotic twins. The values for dizygotic and monozygotic twins were 4 and 17 respectively.

**Conclusion:** The findings of the study show that there is a positive association between sex concordance and breast cancer in female twins though other factors such as zygosity, BMI and age can influence breast cancer diagnosis. From our study, the proband wise concordance for monozygotic twins was higher than that of dizygotic twins. Breast cancer is therefore considered heritable.

## Introduction

Breast cancer remains a major public health problem and its incidence is expected to rise over the next 20years despite various efforts to prevent it (Arnold et al., 2013). This is not surprising because there has been a reported rise in major risk factors of breast cancer in several countries (Howell et al., 2014). The increased risk factors include; obesity, alcohol consumption, inactivity, early menopausal age and late menopausal age. According to Ma & Jemal (2013), breast cancer is the most commonly diagnosed cancer (excluding skin cancer) and the second leading cause of death following lung cancer amongst U.S. women. Sharon et al., (2012) reported that the incidence of breast cancer in men and women is 0.8% and 99.2% respectively.

The influence of exposures at different stages of life as well as genetic factors may increase a person’s risk for breast cancer. These factors act prenatally, in childhood, puberty and reproductively. Many of these are hormonal and/or nutritional, making them very difficult to measure (Howell et al., 2014). Few researches have been done to ascertain the effect of twin birth on breast cancer. It has been shown that, there is an association between circulating androgens and risk of breast cancer (Hui - Chen, 2011).

Women with BRCA1 and BRCA2 mutation have about 87% lifetime risk of developing breast cancer (Palmet & Writer, 2004). Hamilton & Mack (2000), followed identical twins prospectively to find new cases of breast cancer. They found that, cancer incidence was much higher in twins whose mothers had been diagnosed of breast cancer than twins who whose mothers do not have breast cancer. Genetic effects will be important if a twin study shows likelihood of cancer is higher in monozygotic twins (who share all genes) than among dizygotic twins (who, on average, share 50 percent of their segregating genes). However, if the concordance is analogous in both types of twins, shared environmental effects are probably more important (Lichtenstein et al., 2000).

Some studies have been done on the association between intra-uterine variables and risk of breast cancer. For instance, the association between birth weight discordance in twins and risk of breast cancer has been done in the following studies. A case control study in Sweden by Ekbom et al. (1992) showed a positive association between twin birth weight and breast cancer. There was also a positive correlation shown between twin birth weight and risk of breast cancer before 50 years, in a population based, case – control study in Washington (Sanderson et al., (2000). Zech and his colleagues in 2008, reported some relationships between sex –discordance and certain genetic diseases. Hajiebrahimi et al., (2013), in a cross-sectional study reported risk of breast cancer in opposite-sexed twins. Nevertheless, to our best of knowledge, just a few published studies have been looking into sex discordance in twins and risk of breast cancer.

Does male – female twin pairing increase risk of breast cancer? How common is breast cancer in same sex twins as compared to twins who are discordant for sex? What factors may increase the risk of breast cancer in male – female twin paring. These are some of the questions our research seeks to answer.

## Methodology

### Sample

We analyzed twin data from the Washington State Twin Registry (n= 14,462), and the study was approved by the University of Washington Institutional Review Board. Informed Consent was obtained from all participants. The information used for this research is primarily obtained through the Washington State Twin Registry (WSTR). The WSTR sends a snail mail or email to twins who are identified via Department of Licensing (DOL) or request an invitation through the registry’s website. Interested twin pairs are invited and once they consent to participate, they are enrolled into the registry. The study uses a cross-sectional design. Contained in the survey are items on sociodemographic characteristics, habits, healthcare use, physician diagnosed health conditions, standard psychiatric measures and measures of physical health.

### Primary measures

Variables collected include breast cancer (dependent variable), sex pairing (independent variable) sociodemographic characteristics of twins in this study reporting their gender, age, weight, zygosity, sex concordance, age apart and Body Mass Index. Breast cancer was determined by asking questions like “have you ever been diagnosed of breast cancer? Sex Discordance (SD) in twins is defined male –female twin pairing. SD was determined by asking the twin the sex of the other twin.

### Statistical Analysis

Analysis of data was done using the IBM SPSS Statistics for Windows, version 24.0. Data was checked the data for errors, duplicate entries, and retrievable missing data. Data clean up analysis was performed twice via checking frequencies to ensure the validity of data. In cases where the error was found, verification was made by referring to the questionnaires.

The safety of the data was guaranteed by using coding system, making backups and keeping the data in a safe place. We conducted a descriptive analysis of all the variables. An independent T-test was done for continuous variables such as age, age apart and BMI. Chi- square test was used to determine relationship between variables. The primary effect of interest was the relationship between sex discordance and breast cancer. Other variables such as race, weight, BMI, age, age apart were also included in the analysis.

A Generalized Estimating Equation (GEE) modelling was done to determine the relationship between twin pairs and other variables of interest. GEE takes the cluster nature of twins into account since twin data is paired (Wang, 2011). The primary focus of this is the relationship between sex discordance and risk of breast cancer.

Heritability of breast cancer was calculated using the formula 2d/ (2d+b+c), where d is the number of concordant twins and b, c is the number of discordant twins for breast cancer. Probandwise concordance was calculated for monozygotic (MZ) and dizygotic (DZ) pairs. The variable is considered heritable, when probandwise concordance is higher in MZ twins compared to DZ twins (Jahanfar, Lye, & Krishnarajah, 2011).

## Results

### Analysis

#### Descriptive characteristics

Sample characteristics are shown in table 1. The sample included 14462 twins with mean age and standard deviation of 64(12.16). The study population was only females and white (n= 21352, 93.6%). Majority of the participants were monozygotic twins (n = 12522, 54.9%). The mean age apart of the twins was 4.86. The mean BMI was 26.73kg/m^2^. More than half of the participants were female-female twin pairs (12176, n=53.4). The frequency of participants that were sex discordant is (n=4572, 20%). 338 participants had breast cancer.

**TABLE 1.** 1) Sociodemographic and health characteristics of twins from the University of Washington Twin Registry (n=14, 462)

| **Variables** | **Frequency** | **Percentage** |
| --- | --- | --- |
| **Breast cancer** |  |  |
| Yes | 339 | 1.8 |
| No | 18934 | 98.2 |
| **Sex** |  |  |
| Female | 14462 | 100.0 |
| **Sex pairing** |  |  |
| Female-Female twin | 12176 | 53.4 |
| Male-Female twin | 4572 | 20.0 |
| **Weighed more at birth** |  |  |
| You | 3133 | 48.9 |
| Your co-twin | 32769 | 51.1 |
| **Zygosity** |  |  |
| Dizygotic twins | 10288 | 45.1 |
| Monozygotic twins | 12522 | 54.9 |
| **Race** |  |  |
| White | 21352 | 93.6 |
| Not White | 3605 | 6.4 |
| **Sex concordance** |  |  |
| Sex concordant | 18238 | 80.0 |
| Sex discordant | 4572 | 20.0 |

#### Bivariate Analysis

The study participants were divided into two groups (n=338) and without breast cancer (n=14124). We examined the relationship between breast cancer and sex concordance, age, age apart, BMI, sex pairing and zygosity. We found a statistically significant association between breast cancer, age, BMI, sex pairing and sex (p<0.05) (Table 2). There was no statistically significant relationship between race and breast cancer.. There was a statistically significant relationship between breast cancer and BMI with an almost equal prevalence in breast cancer between participants with BMI below 25 and those with BMI 25 and above. A greater percentage of twins who were discordant for sex (2.4%) had breast cancer than those who were concordant for sex (1.6%). The prevalence of breast cancer was higher in dizygotic twins (2.1%) than monozygotic twins (1.5%).

**TABLE 2.**
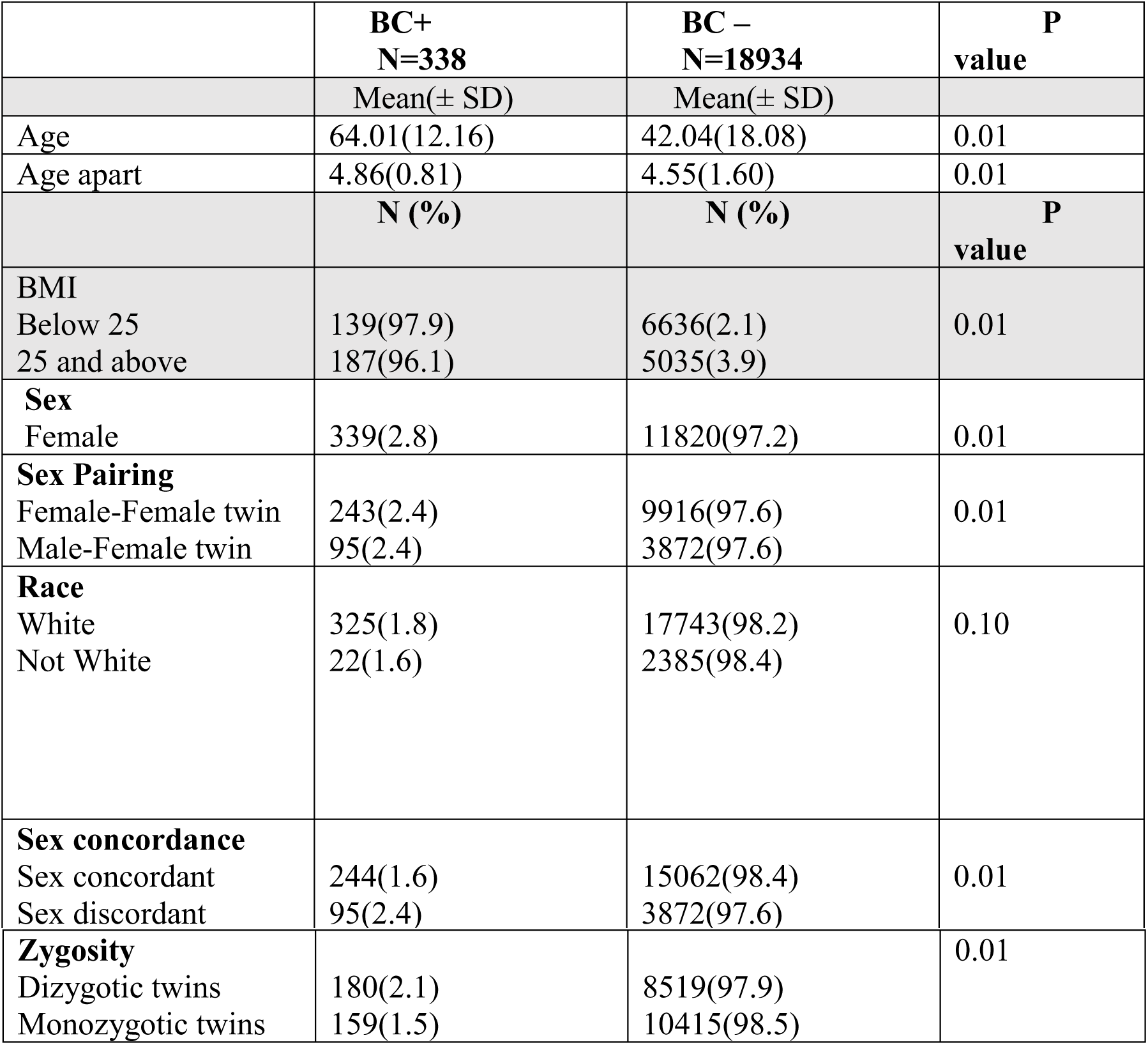
Comparison between twins with and without breast cancer (BC) (n=14, 462)

**Table 3.** Generalized Estimation Equation (GEE)

| VARIABLES | UNADJUSTED ODD RATIO | ADJUSTED ODD RATIO |
| --- | --- | --- |
| **Sex concordance** |  |  |
| Sex concordant | 1.34(1.18-1.53) | 0.69 (0.53-0.90) |
| Sex discordant | 1 | 1 |
| **Zygoty** |  |  |
| Dizygoty twins | 1.20 (1.07-1.34) | 0.78 (0.63-0.97) |
| Monozygoty twins | 1 | 1 |
| **Age** | 0.97(0.96-0.97) | 0.97(0.96-0.97) |
| **Age apart** | 0.94(0.91-0.97) | 1.04(0.96-1.12) |
| **BMI (Kg/m<sup>2</sup>)** |  |  |
| Below 25 | 0.79(0.79-0.88) | 0.94(0.78-1.13) |
| 25 and above | 1 | 1 |
| **Race** |  |  |
| White | 0.78(0.61-1.01) | 0.59(0.29-1.59) |
| Not white | 1 | 1 |
| **Having a child** |  |  |
| Has a child | 0.84(0.69-1.18) | 0.96 (0.76-1.21) |
| Has no child | 1 | 1 |

**Table 4.**
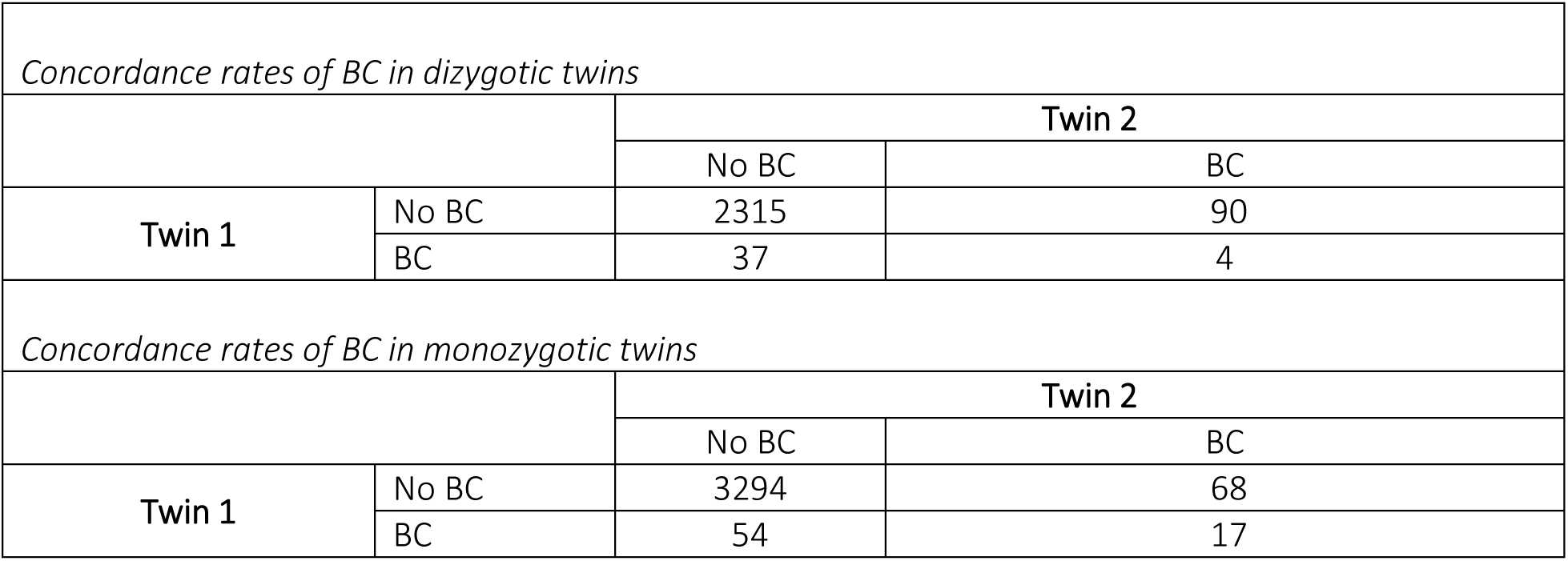

#### Generalized Estimating Equation (Gee)

A GEE analysis was done to predict the odds of breast cancer in female–female twin pairs and male-female twin pairs. Being a female-female twin pair increased the odds of breast cancer by 34% (95%CI: 1.18-1.53). After adjusting for zygosity, age, BMI, race, and childbirth, the odds of breast decreased by 31% in female-female twin pairs [AOR (95%CI):0.69 (0.53-0.90)]. The odds of breast cancer increased by 20% in dizygotic twins as compared to monozygotic twins (95%CI: 1.07-1.34). BMI below 25kg/m^2^, reduced the odds of breast cancer to 79% of the odds in BMI of 25kg/m^2^ and above (95% CI: 0.79-0.88). The risk of breast cancer decreased by 3% with decreasing age (95%CI: 0.96-0.97). Having a child reduced the odds of breast cancer by 84% of the odds of not having a child (95%CI: 0.69-1.18).

Adjusting for all other variables, the odds of breast cancer in dizygotic twins reduced to 78% of the odds of breast cancer in monozygotic twins [AOR (95%CI):0.78(0.63-0.97)]. There was no massive change in odds for age, BMI, race and childbirth after adjusting for all other variables.

The National Institute of Health, (2018), defines heritability as the extent to which differences in people’s genes account for differences in their trait. The probandwise concordance was higher in monozygotic twins as compared to dizygotic twins. The values for dizygotic and monozygotic twins were 4 and 17 respectively.

## Discussion

Sex discordance was found to be associated with risk of breast cancer in twins. This finding is in line with a study by Hajiebrahimi et al., (2013), who reported risk of breast cancer in opposite-sexed twins. Key et al. 2002, explained that, there is an association between circulating androgens and risk of breast cancer. When female twins in opposite-sexed twins are prenatally exposed to androgens from their twin brothers, these females may be at risk of cancer (Zeleniuch-Jacquotte et al., 2012) Androgens interact with estrogens through competitive binding, to sex hormone binding globulin, in female twins with male co-twins in utero (Glinianaia, Magnus, & Harris, 2001).

Monozygotic female twins were found to have lower risk of breast cancer than dizygotic twins. This finding is in accordance with a cohort study done in Finland using 13,176 twins obtained from Finland twin registry to figure out breast cancer risk in monozygotic and dizygotic twins. This study found that compared to dizygotic twins, monozygotic twins have decreased risk of breast cancer (Verkasalo, Kaprio, Pukkala, E, & Koskenvuo, 1999). We found that twins with lower BMI were at a lower risk of breast cancer than those with higher BMI. This is in line with findings of Kaijser et al., (2001), who found increased risk of breast cancer in twins with higher BMI. Mansell, Sprague, Berry, Chisholm, & Trentham-Dietz, (2014), explained that this association is seen in women with hormone receptor –positive breast cancer (has receptor for estrogen) but not in women with hormone receptor negative breast cancer (has no receptor for estrogen).

From our study, the probandwise concordance for monozygotic twins was higher than that of dizygotic twins. Breast cancer is therefore considered heritable. This is in accordance with Moller at al., (2016), who analyzed data from Nordic Twin Study of Cancer to estimate the familial concordance and heritability of breast cancer between monozygotic and dizygotic twins. Their findings indicated that familiar factors explain almost half of the variation in liability to develop breast cancer.

### Limitations of the Study

The study demonstrated statistically significant relationship between sex discordance and risk of breast cancer. However, a variety of factors may have imposed limitations on the results. First, the use of self- reported data has the potential of introducing recall bias and social desirability. Second, the use of cross-sectional study design does not allow us to make conclusions of a causal nature.

### Future Study

More studies should be done on how common gene variations (small changes in genes that are not as significant as mutations) may affect breast cancer risk. Also, intra-uterine conditions that predispose twins to cancers should be given more attention.

## Conclusion

This study used data from Washington Twin Registry to figure out the relationship between sex discordance in twins and risk of breast cancer. The study found a positive correlation between the said variables though other variables such as zygosity, BMI and age can influence breast cancer diagnoses.

## References

Arnold M, Karim-Kos HE, Coebergh JW, Byrnes G, Antilla A, Ferlay J, Renehan AG, Forman D, Soerjomataram: Recent trends in incidence of five common cancers in 26 European countries since 1988: Analysis of the European Cancer Observatory. Eur J Cancer 2013.

Ekbom A, Trichopoulos D, Adami HO, Chung-Cheng H, Shou-Jen L.(1992) Evidence of prenatal influences on breast cancer risk. Lancet; 340: 1015–1018.

Giordano SH, Buzdar AU, Hortobagyi GN. Breast Cancer in Men. Ann Intern Med.; 137:678–687. doi: 10.7326/0003-4819-137-8-200210150-00013.

Glinianaia S. V., Magnus P, & Harris J. R., (2001). Tambs K. Is there a consequence for fetal growth of having an unlike-sexed cohabitant in utero? Int J Epidemiol; 27:657–9.

Hajiebrahimi, M., Bahmanyar, S., Oberg, S., Iliadou, A., & Cnattingius, S. (2013). Breast Cancer Risk in Opposite-Sexed Twins: Influence of Birth Weight and Co-Twin Birth Weight. Jnci-Journal Of The National Cancer Institute, 105(23), 1833–1836.

Hamilton, A., & Mack, T. (2003). Puberty and Genetic Susceptibility to Breast Cancer in a Case– Control Study in Twins. The New England Journal of Medicine, 348(23), 2313–2322.

Howell et al.: Risk determination and prevention of breast cancer. Breast Cancer Research 2014 16:446.

Hui-Chen Wu, Esther M. John, Jennifer S. Ferris, Theresa H. Keegan, Wendy K. Chung, Irene Andrulis, Lissette Delgado-Cruzata, Maya Kappil, Karina Gonzalez, Regina M. Santella and Mary Beth Terry, Global DNA methylation levels in girls with and without a family history of breast cancer, Epigenetics, 6, 1, (29), (2011).

Kaijser, M., Lichtenstein, P., Granath, F., Erlandsson, G., Cnattingius, S., & Ekbom, A. (2001). In Utero Exposures and Breast Cancer: A Study of Opposite-Sexed Twins. Journal of the National Cancer Institute,93(1), 60–62.

Lichtenstein P,Granath F, et a., In utero exposures and breast cancer: a study of opposite-sexed twins. J Natl Cancer Inst. 2001; 93(1):60–62.

Ma J., Jemal A. (2013) Breast Cancer Statistics. In: Ahmad A. (eds) Breast Cancer Metastasis and Drug Resistance. Springer, New York, NY

Moller, S, Mucci, LA, Harris, JR, Scheike, T, Holst, K, Halekoh, U, … Hjelmborg, JB. (2016). The Heritability of Breast Cancer among Women in the Nordic Twin Study of Cancer. Cancer Epidemiology, Biomarkers & Prevention : A Publication Of The American Association For Cancer Research, Cosponsored By The American Society Of Preventive Oncology, 25(1), 145–150.

Munsell, M., Sprague, B., Berry, D., Chisholm, G., & Trentham-Dietz, A. (2014). Body Mass Index and Breast Cancer Risk According to Postmenopausal Estrogen-Progestin Use and Hormone Receptor Status. Epidemiologic Reviews, 36(1), 114–136.

National Institute of Health, (2018). Your Guide to Understanding Genetic Conditions. National Library of Medicine

Parmet, S., Lynm, C., & Glass, R. (2004). Genetics and Breast Cancer. JAMA, 292(4), 522.

Sanderson M, Williams MA, Malone KE, Stanford JL, Emanuel I, White E, Daling JR. (2000). Perinatal factors and risk of breast cancer. Epidemiology;7:34–37.

Southey, M., Joo, J., Dowty, J., Milne, R., Wong, E., Dugue, P., Giles, G. (2017). Heritable methylation marks associated with breast cancer risk. Cancer Research, 77(S13), Cancer Research, 2017 Jul, Vol.77 Suppl 13.

Verkasalo, P., Kaprio, J., Pukkala, E., & Koskenvuo, M. (1999). Breast cancer risk in monozygotic and dizygotic female twins: A 20-year population-based cohort study in Finland from 1976 to 1995. Cancer Epidemiology, Biomarkers & Prevention : A Publication of the American Association for Cancer Research, Cosponsored by the American Society of Preventive Oncology,8(3), 271–4.

Wang, L. (2011). GEE analysis of clustered binary data with diverging number of covariates. ArXiv.org, 39(1), 389–417.

Zech NH, Wisser J, Natalucci G, Riegel M, Baumer A, Schinzel A. Monochorionic–diamniotic twins discordant in gender from a naturally conceived pregnancy through postzygotic sex chromosome loss in a 47,XXY zygote. Prenat Diagn 2008; 28:759–763

Zeleniuch-Jacquotte A, Afanasyeva Y, Kaaks R, et al. Premenopausal serum androgens and breast cancer risk: a nested case-control study. Breast Cancer Res. 2012;14(1):R32

